# How rat flooding occurs during mast seeding of *Melocanna baccifera*?

**DOI:** 10.1101/2025.07.17.665385

**Authors:** Ajikumaran Nair S, Anil John Johnson, Balaji Govindan, Bhaskaran Gopakumar, Konnath Chacko Koshy, Sabulal Baby

## Abstract

Gregarious flowering and mast seeding of bamboos have been linked to rat population outbreaks across multiple geographic regions. This study explores the reproductive mechanism driving the surge in rat population triggered by the fruiting of the bamboo species, *Melocanna baccifera.* We examined four key reproductive parameters, *viz.*, mounting frequency, serum nitric oxide (NO), follicle stimulating hormone (FSH), luteinizing hormone (LH), in mice and rats under *M. baccifera* fruit- or its amino acid (AA)-fed conditions.

The mounting frequency of *M. baccifera* fruit liquid administered male mice was 5.50 ± 1.73, compared to 2.00 ± 0.82 of control animals. NO level in *M. baccifera* fruit liquid-fed mice displayed an increase of 131.83%. The pregnancy and pups number in *Muli* fruit-fed female animals enhanced by 66.66% and 131.25%, respectively. The average pups production in *Muli* fruit-, dAA (d-Thr-Ser-Glu-Phe-His-Ala)-fed and control female rats were 7.40 ± 0.83, 8.80 ± 0.75, and 5.34 ± 0.52 (pups/animal), respectively. *M. baccifera* fruit and dAA combination (dSer-dThr-dGlu) showed increments in FSH and LH levels upto (65%, 58.59%), and (286.41%, 99.24%), respectively.

*Synthesis.* The increased mounting frequency, along with elevated levels of serum NO, FSH, and LH, in animals consuming *M. baccifera* fruits or the AA mix, strongly indicates enhanced reproductive activity. This is the first study providing a mechanistic insight into the rapid rat population boom associated with the cyclical flowering and fruiting of *M. baccifera*. Our findings suggest that the amino acids present in *M. baccifera* fruits may have potential applications in the treatment of infertility.

## 1. INTRODUCTION

*Melocanna baccifera* (Roxb.) Kurz is an economically important bamboo (subfamily Bambusoideae of family Poaceae) well-known for its largest (c. 300 g) baccoid fruits (Banik 2016), and the ecological mayhems associated with its flowering (Govindan et al., 2016; Govindan et al., 2019; Koshy et al., 2022). Natively distributed in the Indian subcontinent to Myanmar it is cultivated in several countries (Vorontsova et al., 2016), and its germplasm has been conserved in 14 bambuseta in six countries (Koshy, 2023; Koshy and Baby, 2025). The young shoots, culms, leaves and fruits of this bamboo are used in various ways such as food, medicine, paper and traditional industries (Koshy and Baby, 2025). A series of previous studies in *M. baccifera*, highlighted its phytochemistry, nutritional aspects, flower-fruit dynamics, and visitor-predator patterns (Govindan et al., 2016, Govindan et al., 2019; Koshy et al., 2022).

Gregarious flowering and mast seeding of *M. baccifera* occurs every 40 to 50 years (Gamble, 1896; McClure, 1966; Janzen, 1976, Ramanayake and Weerawardene, 2003; Banik, 2010; Koshy et al., 2022), and each clump produces large quantities of fleshy fruits (37.19 to 456.67 Kg/clump), averaging 4796.38 (± 3420.26, n = 8) fruits, weighing 237.18 (± 169.13 Kg, n = 8) (Koshy et al., 2022). Though an array of animals predates on these fruits, the intense predation by rats results in sudden increase in their numbers (rat flooding). When the quantity of bamboo fruits diminishes, rats attack other grain crops ultimately leading to food shortage, famines, human loss, and even political uprisings (John and Nadgauda, 2002; Jeeva et al., 2009; Aplin and Lalsiamliana, 2010; Belmain et al., 2010; Htwe et al., 2010; Ruiz-Sanchez and Sosa, 2015; Govindan et al., 2016; Koshy et al., 2022). Mizoram, one of the northeastern states in India, is the epicenter of this ecological phenomenon, locally known as *Mautam* (Gamble, 1896; McClure, 1966; Janzen, 1976; Ramanayake and Weerawardene, 2003; Banik, 2010; Koshy et al., 2022). Due to the resultant ecological and humanitarian crisis in relation to its mast seeding, previous flowering incidences of this bamboo gained extensive media attention (Shoumatoff, 2008; Normile, 2010; Wilkins, 2010).

Mizoram is home to vast bamboo forests covering approx. 6,447 sq. km, around 31% of the State’s total geographical area. Mizoram harbors over 20 bamboo species, of which *M. baccifera*, locally known as *Muli*, constitutes more than 95% of the total bamboo stock. To mitigate the impact of the latest *M. baccifera* gregarious flowering (2005-2009) and the ensuing rodent outbreaks (Aplin and Lalsiamliana, 2010), the State government launched the Bamboo Flowering and Famine Combat Scheme (BAFFACOS) in August 2004 (BAFFACOS, 2009). Under this scheme, farmers were encouraged to control rat populations using traditional traps and chemical rodenticides, which were distributed free of cost. Curiously, despite bounty payments totaling Indian Rupees 29.65 lakhs (34,398.13 USD) for 1.51 million rat tails collected in 2006-07, rats still caused considerable damage to jhum paddy, vegetables, fruits, and lowland rice cultivation affecting nearly 82.9% of the State’s total cropping area in 2007-08 (BAFFACOS, 2009). Several accounts of *Mautam* in Mizoram highlight a surge in the litter size of rats feeding on bamboo fruit, with some even narrating rats as large as wild cats (examples: Shoumatoff, 2008; Aplin and Lalsiamliana, 2010). These rat tales and statistics of *Mautam* highlight the need for a deeper understanding of this ecological phenomenon.

Previous studies have reported the presence of various bioactive constituents in *Melocanna baccifera* fruits, such as amino acids, sugars, phenolic acids, fatty acids, vitamin B3, and minerals, indicating their potential nutritional and therapeutic value. The protein content in its fruits is notably low. *In vivo* rat feeding experiments demonstrated that *M. baccifera* fruit alone is insufficient to support the normal growth and physiological functions in rats. However, when combined with a normal diet, bamboo fruit helped maintain the body weight, serum biochemical and hematological parameters within normal ranges, along with a reduction in serum total cholesterol levels (Govindan et al., 2016). Fruit production gradually increases from a nominal level in the first year of seeding, peaks in the second or third year and then declines, ultimately leading to the death of the clumps. Previous studies perceived a link between the ecology of *M. baccifera* (fruits) and its chemistry based on the exciting interactions of visitors and predators observed during mast seeding, and provided preliminary leads towards explaining the cause of rat multiplication (Govindan et al., 2016; Koshy et al., 2022).

Here we elucidate the molecular mechanism behind the rise in rat population due to *M. baccifera* fruiting, which leads to *Mautam*.

## 2. MATERIALS AND METHODS

### 2.1. Melocanna baccifera fruits

The study was undertaken on *Melocanna baccifera* clumps situated in the Bambusetum of the Jawaharlal Nehru Tropical Botanic Garden and Research Institute (JNTBGRI) at Palode in Kerala, southern India (Koshy, 2010; Koshy, 2017; Koshy et al., 2022); gregarious flowering of these bamboo clumps during 2009 to 2015 produced flowers and fruits. Flowering in clump 404 was initiated in September 2012, and is still being continued (Koshy et al., 2022). *M. baccifera* fruits for our studies were gathered from eight clumps with accession numbers 58, 359, 365, 394, 395, 403, 404, 405.

### 2.2. Animals

Inbred male and female Wistar rats (*Rattus norvegicus*) and Swiss albino mice (*Mus musculus*), housed under uniform hygienic conditions in the Institute Animal House and fed with standard pellet feed (PF, Kamadhenu Feeds, Bangalore) and water *ad libitum*, were used for the experiments. Animals were kept under a 12-hour light/dark cycle, at a temperature of 23-25°C and relative humidity between 45% and 55% throughout the experimental period. These animal experiments were approved by The Institute Animal Ethics Committee (IAEC) (B-01/12/2011/PPD-03 dated 19.12.2011 & B1/02/2023/PPD-01 dated 03.11.2023). Animal studies were conducted in accordance with the guidelines set by the Committee for the Control and Supervision of Experiments on Animals (CCSEA), Government of India.

### 2.3. Mounting frequency

The mounting frequency in mice was determined as described in Subramoniam and coworkers (1997). Briefly, twelve adult male Swiss albino mice (26-29 g body weight) were divided into two groups of six animals each, along with two similar groups of age and weight matched, adult non-estrus female mice separately. The male mice in the control and test groups were administered with a single per oral dose (0.5 mL/animal) of water or *M. baccifera* fruit liquid (21-28 days old fruit), respectively, and 30 min after the administration of water or fruit liquid, the male mice were individually placed in a clean aquarium and stayed acclimatized for 15 min. After acclimatization the males were paired individually with non-estrus female mouse for 3 h. Number of mounts in both control and test animals were recorded by two observers in 15 min at the start of every 1 h interval for 3 hrs (Fig. 1). All experiments were performed at 27-29°C room temperature from 09.00 am to 12.00 noon on sunny days.

**Figure 1.**
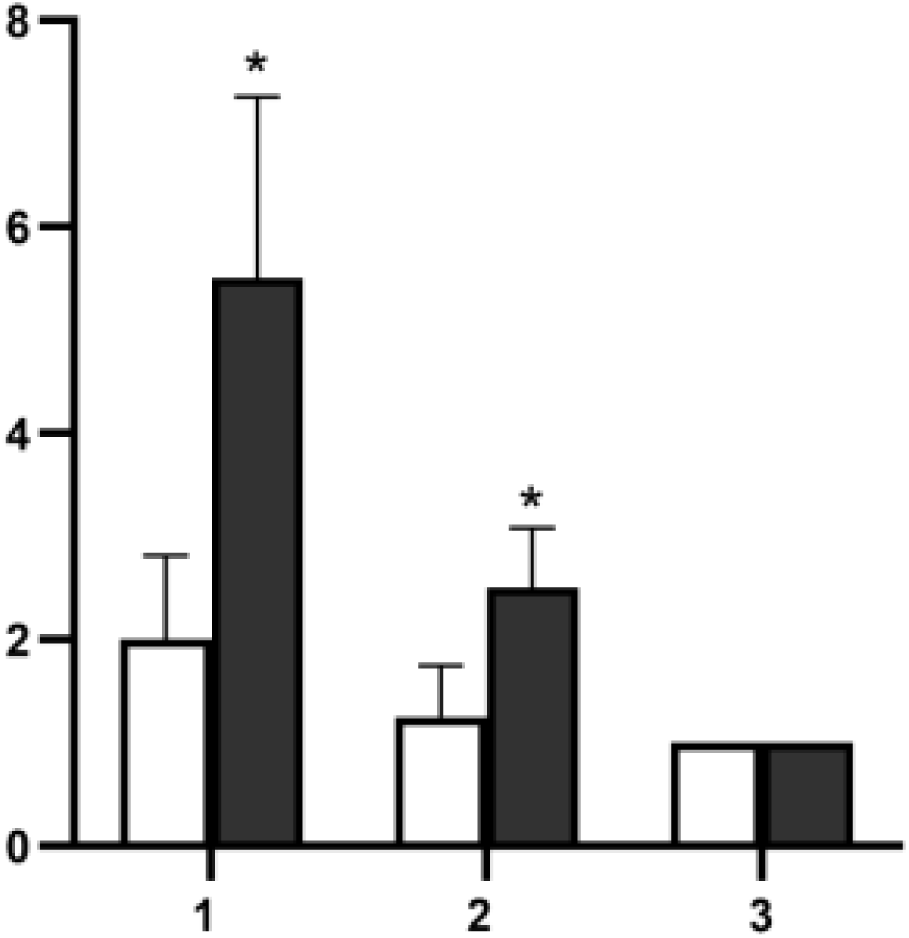
Mounting frequency in male mice fed with water or bamboo fruit liquid (filled bars) in 3 hrs. Values are mean ± SD, n = 6, *p ≤ 0.05 (compared to control, unfilled bars).

### 2.4. Serum nitric oxide level in mice

To assess nitric oxide (NO) levels in the blood, twelve adult male Swiss albino mice (22-25 g body weight) were randomly divided into two groups of six animals each. Animals in control and test groups were administered with a single per oral dose (0.5 mL/animal) of distilled water or *M. baccifera* fruit liquid (21-28 days old fruit), respectively. Animals were sacrificed by cervical dislocation 30 min after administration of either water or fruit liquid. Blood samples were collected *via* cardiac puncture, and serum NO levels were measured immediately (Sastry et al., 2002).

### 2.5. Nitric oxide level in mice blood

Serum NO levels in control or test animals were determined indirectly by estimating the serum nitrate/nitrite levels as described by Sastry and co-workers (2002). Briefly, 100 μL serum samples of each control or test animals were added with 400 μL of carbonate buffer (50 mM, pH 9.0), and mixed well with small amount of freshly activated cadmium granules (0.20 g). The mixtures in test tubes were incubated at room temperature for 1 h with thorough shaking. The reaction was terminated by adding NaOH (35 mM, 100 μL) and ZnSO₄ (120 mM, 400 μL) under vortex, followed by a 10 min incubation. The tubes were then centrifuged at 4000 rpm and supernatants were separated. Supernatant (500 μL) from each control or test or standard were then added with 1.0% sulfanilamide (250 μL, in 3 N HCl) and 1.0% N-1-naphthylethylenediamine (250 μL, in water) and mixed well. After 10 min, optical densities were measured at 540 nm in a UV1650PC spectrophotometer (Shimadzu, Japan). Along with the serum samples, standards and blank groups were worked out. In standard group instead of serum, nitrite standards were added and in blank no standard or serum was added. Standard graph was plotted and serum NO levels in the control and test groups were determined.

### 2.6. *Muli* fruits, amino acids and rat reproductive performance

Male and female Wistar rats of same age groups (7 weeks) with average body weight of 180-200 g were selected for the experiments. In order to test the effect of bamboo fruit on female reproductive capacity (rat pup production), female rats were grouped into control, bamboo fruit test and amino acid (AA) test animals. In these groups, animals were kept in polypropylene cages in uniform manner with 2 animals/cage. Female rats in (i) control group were provided daily with pellet feed (PF, 200 g/cage) and water (W, 100 ml/cage), (ii) bamboo fruit (BF) group were provided with mature (42 days age) bamboo fruit pericarp (100 g/cage) + pellet feed (100 g/cage) and water (100 ml/cage), and (iii) in AA group were provided with pellet feed (200 g/cage) and water (100 ml per cage) dissolved with 0.01% of six major AAs (d-threonine (d-Thr), d-serine (d-Ser), d-glutamic acid (d-Glu), d-alanine (d-Ala), d-phenylalanine (d-Phe), d-histidine (d-His)) in *Muli* fruit liquid (35 days age) and mature fruit pericarp (42 days age) or their combinations (Govindan et al., 2016). In three AA combinations, the six AAs (Thr, Ser, Glu, Ala, Phe, His) were divided in two sets of three AAs each (Thr, Ser, Glu & Ala, Phe, His), and their 0.01% combinations were dissolved in water (Table 1). These two combinations of AAs (d or l AAs; d-Thr, Ser, Glu or l-Thr, Ser, Glu, or d-Ala, Phe, His or l-Ala, Phe, His) were provided to female rats (four groups) along with (PF+W)-fed control groups. All animals were provided with pellet feed. Animals in these experiments were free to access pellet feed or test materials or water as much as they desired. In every day, after 24 hrs, the PF, BF, AA-water and water were replaced with fresh ones after measuring the daily consumption, and the feeding was continued for seven days.

**Table 1.**
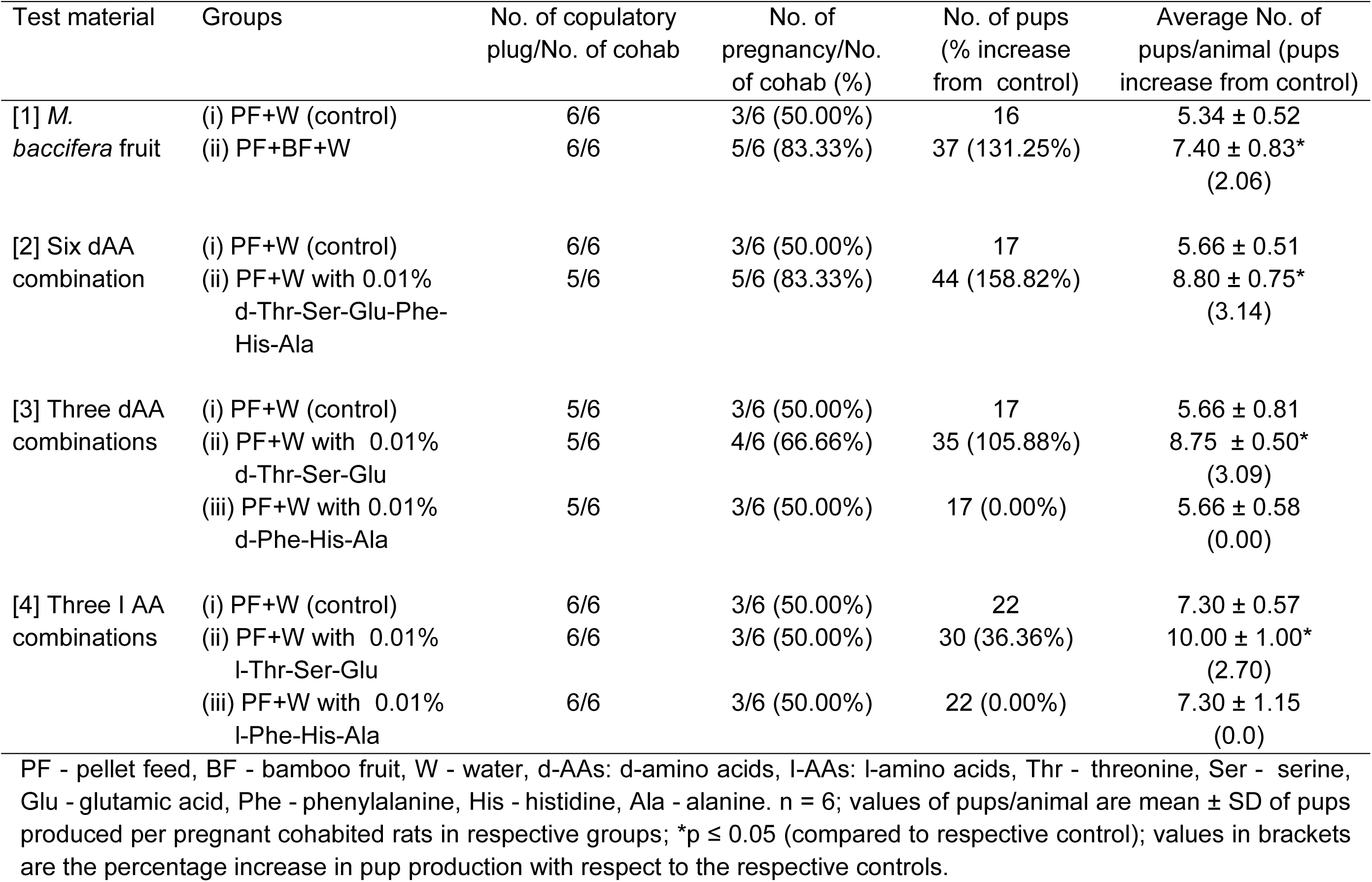
Reproductive performance in female rats fed with *M. baccifera* fruit and amino acid combinations.

| Test material | Groups | No. of copulatory plug/No. of cohab | No. of pregnancy/No. of cohab (%) | No. of pups (% increase from control) | Average No. of pups/animal (pups increase from control) |
| --- | --- | --- | --- | --- | --- |
| [1] *M. baccifera* fruit | (i) PF+W (control) | 6/6 | 3/6 (50.00%) | 16 | 5.34 ± 0.52 |
|  | (ii) PF+BF+W | 6/6 | 5/6 (83.33%) | 37 (131.25%) | 7.40 ± 0.83*<br>(2.06) |
| [2] Six dAA combination | (i) PF+W (control) | 6/6 | 3/6 (50.00%) | 17 | 5.66 ± 0.51 |
|  | (ii) PF+W with 0.01% d-Thr-Ser-Glu-Phe-His-Ala | 5/6 | 5/6 (83.33%) | 44 (158.82%) | 8.80 ± 0.75*<br>(3.14) |
| [3] Three dAA combinations | (i) PF+W (control) | 5/6 | 3/6 (50.00%) | 17 | 5.66 ± 0.81 |
|  | (ii) PF+W with 0.01% d-Thr-Ser-Glu | 5/6 | 4/6 (66.66%) | 35 (105.88%) | 8.75 ± 0.50*<br>(3.09) |
|  | (iii) PF+W with 0.01% d-Phe-His-Ala | 5/6 | 3/6 (50.00%) | 17 (0.00%) | 5.66 ± 0.58<br>(0.00) |
| [4] Three l AA combinations | (i) PF+W (control) | 6/6 | 3/6 (50.00%) | 22 | 7.30 ± 0.57 |
|  | (ii) PF+W with 0.01% l-Thr-Ser-Glu | 6/6 | 3/6 (50.00%) | 30 (36.36%) | 10.00 ± 1.00*<br>(2.70) |
|  | (iii) PF+W with 0.01% l-Phe-His-Ala | 6/6 | 3/6 (50.00%) | 22 (0.00%) | 7.30 ± 1.15<br>(0.0) |

PF - pellet feed, BF - bamboo fruit, W - water, d-AAAs: d-amino acids, l-AAAs: l-amino acids, Thr - threonine, Ser - serine, Glu - glutamic acid, Phe - phenylalanine, His - histidine, Ala - alanine. n = 6; values of pups/animal are mean ± SD of pups produced per pregnant cohabited rats in respective groups; \*p ≤ 0.05 (compared to respective control); values in brackets are the percentage increase in pup production with respect to the respective controls.

After seven days the female rats in estrous cycles from each group were carefully selected (6 animals/group) and individually cohabited with male rats (one pair/cage) for 48 hrs. After 48 hrs, male animals were removed from cages and the female animals were observed for the copulation plug and maintained with supply of pellet feed and water until their delivery. The number of pups produced was counted for test and control animals. The results of these rat reproduction trials are tabulated in Table 1.

### 2.7. Follicle stimulating, luteinizing hormones in rats

Serum follicle stimulating (FSH) and luteinizing (LH) hormone levels in rat blood were determined as described elsewhere (Diebel and Bogdanove, 1978). Briefly, adult female Wistar rats (182-190 g body weight) with synchronized estrous cycles were divided into three groups, of which the control group received standard rat pellet feed and water; the second group was fed with pellet feed, water, and *Melocanna baccifera* fruit (42-day mature); and the third group received pellet feed and water supplemented with 0.01% d-amino acids (Ser, Thr, Glu). All animals are provided *ad libitum* of water, pellet feed or *M. baccifera* fruit or AAs as specified in respective groups. The feeding was continued for seven days in an identical manner with various feeds (200 g pellet feed and 100 ml water or 100 g pellet feed +100 g *M. baccifera* fruit and 100 ml water or 200 g pellet feed and 100 mL 0.01% d-AAs in100 mL water/two animal/cage/day). The vaginal smear of the animals in three groups was examined to determine their sex cycles. Six animals from each group in the pro-estrus phase were selected and sacrificed by cervical dislocation at 17:00 h, followed by blood collection *via* cardiac puncture.

The blood serum was separated immediately and FSH and LH levels were determined with commercially available ELISA kit following the manufacturer’s instruction manual (Fig. 2).

**Figure 2.**
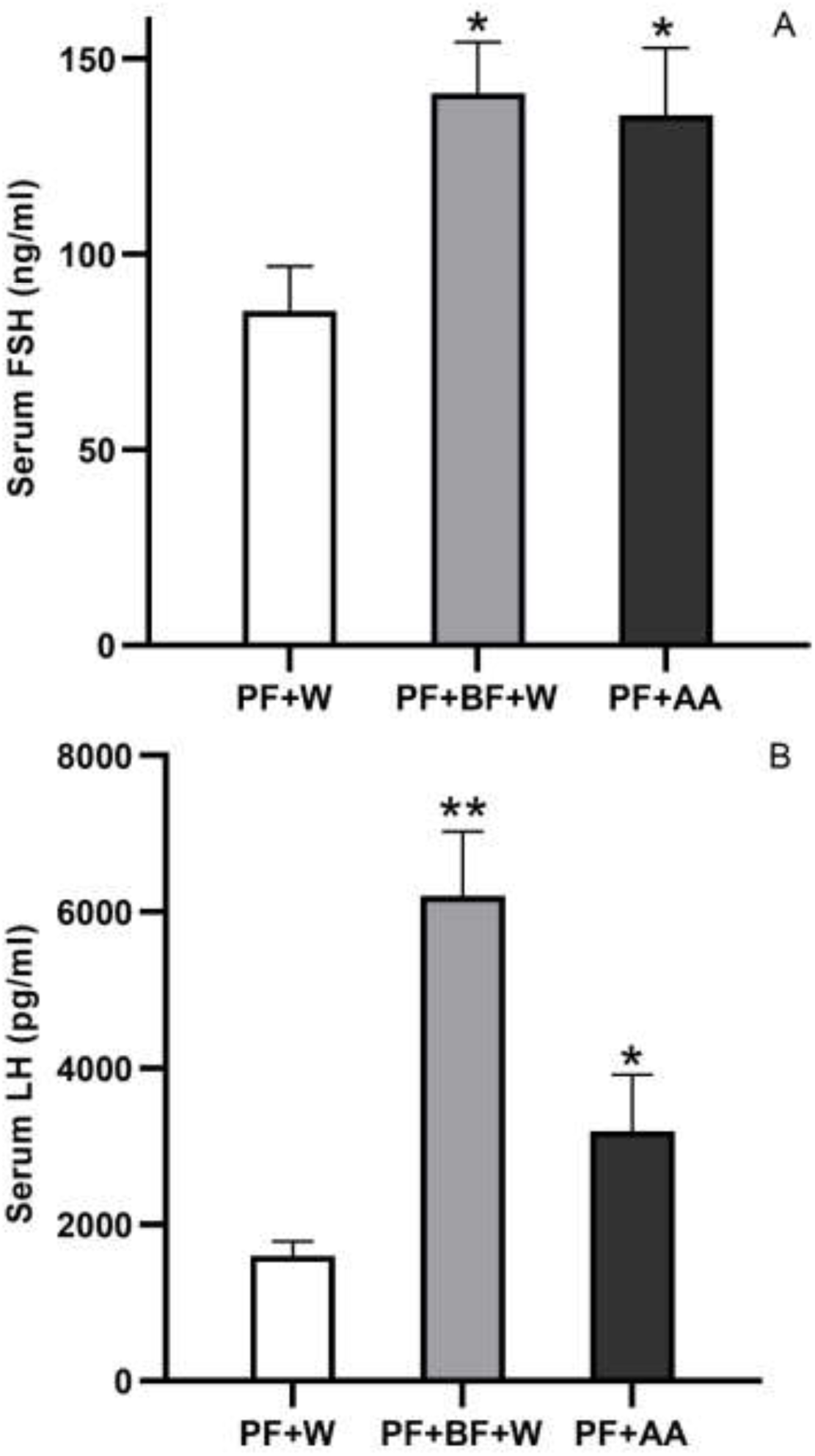
Serum [A] FSH and [B] LH levels. PF+W: rat pellet feed and water; PF+BF+W: rat pellet feed + *M. baccifera* fruit and water; PF+AA rat pellet feed and 0.01% d-AAs (Ser, Thr, Glu) in water; values are mean ± SD, n = 6, **p ≤ 0.01, *p ≤ 0.05 (compared to control, PF+W).

### 2.8. Statistical analysis

Results from various assays are presented as mean ± standard deviation (S.D.). Statistical comparisons were performed using one-way ANOVA followed by Dunnett’s post-hoc test. For comparisons involving only two groups, an unpaired Student’s t-test was applied. A *p*-value of less than 0.05 was considered statistically significant.

## 3. RESULTS

### 3.1. Mounting frequency, serum nitric oxide

The mounting frequency after 1 h for *M. baccifera* fruit liquid administered male mice was 5.50 ± 1.73 in comparison to 2.00 ± 0.82 to water-fed control animals. After 2 hrs the mounting frequencies in fruit liquid administered and control male mice were 2.5 ± 0.58 and 1.25 ± 0.50, respectively. After third hour, the control and test mice did not show any significant difference in mounting frequencies and the values in both the cases were 1.0 ± 0.00 (Fig. 1). Control mice which received 0.5 ml of water showed an average serum NO level of 43.48 ± 5.97 μmol/L, and the same dose of *M. baccifera* fruit liquid showed a significant increase in the serum NO level in test animals (100.8 ± 18.05 μmol/L). The increment in NO level in test mice is 131.83% than that of the control animals.

### 3.2. *Muli* fruits, amino acids and rat reproductive performance

In rat reproductive performance trials influenced by *Muli* fruits, six d-AA, three d or l-AA combinations, female rats fed with (PF+W) or (PF+BF+W) individually cohabited with male rats on estrus stage showed successful copulation in 48 hrs of mating period, and displayed 100% copulation plug in both (PF+W) and (PF+BF+W)-fed female animals. Out of these 6 animals each in the control and test groups, 3 control and 5 test animals displayed successful pregnancy, respectively. The increase in pregnancy in *Muli* fruit-fed female animals was 66.66% than that of the control animals. The litter size also increased in *Muli* fruits fed rats, wherein 131.25% increase in number of pups was observed. The average number of pups produced per animal was 7.40 ± 0.83 in *Muli* fruit-fed rats and 5.34 ± 0.52 (only) in control rats.

In PF + six dAA (0.01% in water)-fed female rats the pregnancy rate is 83.33%, but in its control rats the pregnancy rate is only 50.00%; these test animals displayed a 66.6% increase in pregnancy rate compared to the control. Again, in PF + six dAA-fed female rats the litter size displayed 158.82% increase from the control animals fed with rat pellet feed and water. The average pups production/rats in PF + six dAA-fed female rats were 8.80 ± 0.75 (pups/animal), and the value in control animal was 5.66 ± 0.51 (pups/animal) (Table 1). PF + six dAA-fed female rats displayed significant increase in pups production to that of the control animals and the increase in number was 3.14 pups/animal. Among the female rats fed with PF + 3 dAAs (0.01% in water) combinations, PF + 0.01% d-Thr-Ser-Glu in water-fed group displayed significant increase in pregnancy rate as well as pups production (compared to control animals), and the values correspond to 33.33% and 105.88%, respectively. The average pups/animal in PF + 0.01% d-Thr-Ser-Glu in water-fed animals were 8.75 ± 0.50 with an increase of 3.09 pups/animals from the control group. PF+ 0.01% d-Phe-His-Ala in water-fed group didn’t show any increase in these parameters from the control animals (Table 1).

The pregnancy rate in PF+W, PF+0.01% l-Thr-Ser-Glu in water and PF+0.01% l-Phe-His-Ala in water fed rats were same (50.00%), and the female rat fed with PF+ 0.01% l-Thr-Ser-Glu in water only displayed 36.36% increase in pregnancy rate to that of control animals. The average pups production/animals in PF+W, PF+ 0.01% l-Thr-Ser-Glu in water and PF+ 0.01% l-Phe-His-Ala in water fed rats were 7.30 ± 0.57, 10.00 ± 1.00 and 7.30 ± 1.15, respectively. Among which PF+ 0.01% l-Thr-Ser-Glu in water fed animals exhibited an increase of 2.70 pups/animal to that of control rats. The multiplication rates with individual AAs d or l (Phe-His-Ala) were not as promising as the combination of three d or l AAs (Thr-Ser-Glu) (Table 1).

In rat reproductive performance experiments, average weight gain for female control animals (4 groups, 24 animals, just before pairing) was +12.6 ± 7.47 g (n = 24).

Similarly, average weight gain for test female animals (6 groups, 36 animals, just before pairing) was +20.51 ± 11.60 g (n = 36). The average of food-water intakes in test rats (2 animals per cage, seven days collective average): pellet food : bamboo fruit: (bamboo fruit, after subtracting max. wt. reduction):water = 25.21 ± 8.22 (g):32.51 ± 8.08 (g):(18.28 ± 5.69 g):40.75 ± 10.61 (ml) (n = 36)). Average of food and water intake in control rats (2 animals per cage, nine days) were 25.22 ± 7.61 g and 47.27 ± 11.20 mL respectively (n = 24). Eight and ninth days (after pairing male with female) the overall food/water intake reduced a little due to increased mounting activity evidenced by the copulation plugs in female rats. Average normal food intakes in both test and control rats in all groups were nearly the same. In (PF+BF+W)-fed test animals, in addition to an equal amount of normal food consumption the female rats consumed an average of 32.51 ± 8.08 g of bamboo fruit/animal. Even after subtracting a maximum weight reduction of bamboo fruit in 24 hrs under identical animal house conditions the fruit intake in (PF+BF+W)-fed animals were 18.28 ± 5.69 g/animal. In this test group water intake was slightly reduced to 40.75 ± 10.61 mL compared to control animals 47.27 ± 11.20 mL, and this lesser intake of water accounts for the water content in consumed *Muli* fruits in (PF+BF+W)-fed test animals.

### 3.3. Follicle-stimulating, luteinizing hormones in rats

Serum FSH values in the control (PF+W)-fed animals were 85.55 ± 11.5 ng/mL, and that of (PF+BF+W)-fed animals were 141.22 ± 13.09 ng/mL (Fig. 2A). In (PF+W with 0.01% d-Ser-dThr-dGlu)-fed animals the FSH levels were 135.68 ±17.24 ng/mL. *M. baccifera* fruit and the dAA combination (dSer-dThr-dGlu) showed an increment in FSH to 65% and 58.59%, respectively. Similarly, the control and (PF+BF+W)-fed animals showed serum LH levels of 1605.55 ± 174.70 pg/mL and 6203.54 ± 825.66 pg/mL, respectively, and (PF+W with 0.01% d-Ser-dThr-dGlu)-fed animals displayed 3199.8 ± 716.94 pg/ml of LH. The increment of LH levels in *M. baccifera* fruit and dAA mixture (dSer-dThr-dGlu) treated rats were 286.41% and 99.24%, respectively, than that of control animals (Fig. 2B).

## 4. DISCUSSION

### 4.1. *Muli* fruits, amino acids, rat reproductive performance

Our rat feeding experiments undoubtedly demonstrate the affinity of rats towards fleshy *Muli* fruits. A previous study, through similar feeding experiments, demonstrated the enhanced affinity of rats towards *M. baccifera* fruit liquid and seed (Govindan et al., 2016). Our data indicate that *Muli* fruits are significantly enhancing the reproductive rates in rats, and fruit AAs are the major constituents causing the higher reproductive performance (number of pups and pregnancy rate). A combination of only 3 major d or l-AAs (Thr-Ser-Glu) gave significant results. *Muli* seeds are enriched with sugars, AAs, and other metabolites such as phenolics, alkaloids, steroids, terpenoids, flavonoids, nor-isoprenoids, and fatty acid derivatives (Govindan et al., 2016; Govindan et al., 2019).

The average increase in pups/rat in bamboo fruit fed animals to the control animals were 2.06 pups/animal. This increase in rat pups production by the bamboo fruit intake causes an exponential increase in rat population (Normile, 2010; Wilkins, 2010). The Mizoram government’s BAFFACOS report infers the optimum *M. baccifera* fruit production in the 2nd and 3rd years of fruiting (2006-2007), and the rat flooding also occurred in this period (BAFFACOS, 2009). Fruiting diminished by the end of 3rd to 4th year, which is matching the duration of grain damage. Koshy and co-workers (2022) reported a similar pattern of fruit production and rat predation at JNTBGRI Bambusetum located at the southern end of India. Of the eight clumps studied, three produced the highest fruit yield in the first year, four in the second year, and one in the third year. Fruit production remained high during the first three years, gradually declined thereafter, and culminated in the death of the clumps - except for one (clump 404), which continues to bear fruit even after 12 years and 7 months as of March 2025. Moreover, over 80% of the total fruit production occurred between April and July, coinciding with the peak rat predation in this period (Koshy et al., 2022). Assuming the full-fledged *M. baccifera* fruiting span as three years (approx. 156.54 weeks; Aplin and Lalsiamliana, 2010; Koshy et al., 2022), and considering that rats can reproduce every 8-12 (10) weeks, up to 15.6 generations could occur during this period. Under optimal ecological conditions such as abundant food, no predation and favorable environment, an increase of 2.06 pups per rat attributed to the nutritional boost from bamboo fruit consumption could result in an exponential population growth. Extrapolating these data, a single rat could potentially give rise to approx. 81,662 (0.082 million) descendants over the three year bamboo fruiting span.

### 4.2. Rat multiplication mechanism

The reproductive activity in female rat is influenced by the coordinated action of brain-ovary axis involving GnRH, FSH, LSH, NO and other ovarian hormones. In male rodents it involves the brain-testis axis through androgenic hormones and NO. In *Muli* fruit liquid-fed male mice 131.83% increase in serum NO was observed along with an increase in mounting frequency (5.50 ± 1.73 in 1 h). This co-related action between the bamboo fruit liquid intake and increase in mounting frequency is through the production NO; its production in corpus cavernosa is through the action of endothelial NO synthase (eNOS) enzyme. The phosphorylation of protein kinase B through the phosphorylated phosphatidyl-inositol 3-kinase also plays a pivotal role in the phosphorylation of eNOS to generate NO in penile tissue (Burnett, 2004).

Nitric oxide (NO) is a free radical gas which is considered the main mediator in many cardiovascular, reproductive, digestive and immune system functions. Again, NO is a messenger molecule of neurons, and a vasodilator produced by endothelial cells. The endothelial NO mediates the increase in cGMP through guanylyl cyclase, which intern leads to the relaxation of smooth muscle of the corpora cavernosa culminating in penile erection by vasodilation of the corpora. In fact, NO plays a crucial role in penile physiology, serving as the primary mediator of erectile function (Burnett et al., 1992; Hull et al., 1994; Burnett, 1995; Burnett, 1997; Burnett, 2006; Burnett, 2024). Elevated blood NO levels result in efficient erection and increase in mounting frequency in animals. The increased NO in the corpus cavernosum of the mice fed with bamboo fruit liquid orchestrates the activation of soluble guanylyl cyclase, which intern increases the 3′,5′-cyclic guanosine monophosphate (cGMP) levels, leading to the relaxation of corpus cavernosum smooth muscle culminating in penile erection and increased mounting frequency (Burnett, 2006). Our data demonstrate that the bamboo fruit liquid increases mounting frequency in male rodents through up-regulation of NO activated pathways, leading to increase in rat population (Fig. 1).

Female rats fed with (PF+BF+W) or (6 dAAs) or (3 d/l AAs: Ser-Thr-Glu) combination displayed significant increase in production of pups (Table 1), and the female rats fed with these feed combinations also displayed significant hike in FSH and LSH, the key hormones involved in reproduction. The brain-ovary axis influenced by FSH and LSH leads to the increase in successful pregnancy and pups production. The BF and the tested AAs (3 dAAs, Thr-Ser-Glu) displayed elevated FSH/LSH production from pituitary gland in female rats due to elevated or prolonged action of gonadotropin-releasing hormone (GnRH) raised from the hypothalamus. GnRH is a deca-peptide which comprises of a highly conserved ten AA sequence (pGlu-His-Trp-Ser-Tyr-Gly-Leu-Arg-Pro-Gly-NH2), of which the AAs Ser and Glu are present in the tested three AA combination. *Muli* fruit pericarp and fruit liquid are rich sources of AAs (Govindan et al., 2016), and consumption these fruits by rats as food produces GnRH constantly in an orchestrated manner leading to the elevated production of FSH/LSH, resulting in increase in pups production. It is also a proven fact that the essential AAs are a prerequisite for young rats in attaining timely reproductive maturity (Liu et al., 2024).

The AA residues flanking Glu in GnRH decapeptide sequence displayed crucial role in selectivity and interaction in mammalian and non-mammalian GnRHRs (Wang et al., 2008). In our study, we found that the bamboo fruit fed rodents produce NO in elevated condition, which is in correlation with the findings of Clarkson and Herbison (2006). It was reported that the preovulatory surge of GnRH is crucial for reproduction in mammals, and the neurotransmitters glutamate and NO play major roles in reproduction. GABA_A_ (AA neurotransmitter) receptor signaling develops in advance of glutamatergic signaling. But, the switch from GABA_A_ receptor depolarization to hyperpolarization of GnRH neurons is delayed until puberty. These findings suggest that developing GnRH neurons exhibit a sequence of GABA-glutamate signaling (similar to other neuronal networks), but it is significantly elongated so as to be complete only by the time of puberty onset. In this study, one of the crucial AAs is Glu in the bamboo fruits. Again, studies have shown that elevated glutamate levels in several hypothalamic nuclei play a key role in GnRH release (Brann, 1995; Iremonger et al., 2010). Glutamate receptors are also located in these key hypothalamic nuclei. Gonadotropins stimulate the gonads in males and the ovaries in females.

In females, an acute rise of LH triggers ovulation and development of the corpus luteum. In males, LH stimulates the production of testosterone by the Leydig cells. It acts synergistically with FSH, which stimulates the ovaries to grow ovarian follicle, causing the follicle cells to secrete estrogen. LH causes ovulation and follicle cells of the corpus luteum to secrete progesterone. So, FSH and LH control the release of estrogen and progesterone, respectively (Scheme 1). Glutamate agonists are known to stimulate GnRH and LH release, and glutamate receptor antagonists attenuate the steroid-induced and preovulatory LH surge. Glutamate is also involved in the critical processes of puberty, hormone pulsatility and sexual behavior. Glutamate elicits these effects by activating the release of the gaseous neurotransmitter, NO, which potently stimulates GnRH (Dhandapani and Brann, 2000). Barnes and co-workers showed that chronic NO deficiency is associated with a decrease GnRH release, which subsequently leads to a significant decrease in the amount of FSH- and LH-stimulated release from the pituitary. Otherwise, the release of FSH and LH is not only under the control of GnRH, but is also controlled by NO (Barnes et al., 2001). Several other studies resulted in similar findings (Iremonger et al., 2010; Farkas et al., 2018). Estradiol (estrogen steroid hormone) is the major female sex hormone. It is involved in the regulation of the estrous and menstrual female reproductive cycles. Recently, Farkas and co-workers showed that estradiol facilitated neurotransmissions to GnRH neurons *via* both GABA_A_-R and glutamate/AMPA/kainate-R. Through the ERβ/Akt/nNOS pathway, estradiol acts directly on GnRH neurons in proestrus-afternoon through generating NO, which retrogradely accelerates GABA and glutamate release from the presynaptic terminals contacting GnRH neurons. This mechanism might be contributing to the regulation of the GnRH surge, a fundamental prerequisite of ovulation (Farkas et al., 2018) (Scheme 1). Briefly, the stimulation of the reproductive axis in rats fed with *B. baccifera* fruits involves the key regulators GnRH, GABA, and NO, which leads to increased secretion of FSH and LH, and this hormonal surge enhances increase in ovulation, implantation and maintenance of fetus and successful delivery of higher number of pups in *Muli* fruit- or AA-fed female rats.

**Scheme 1.**
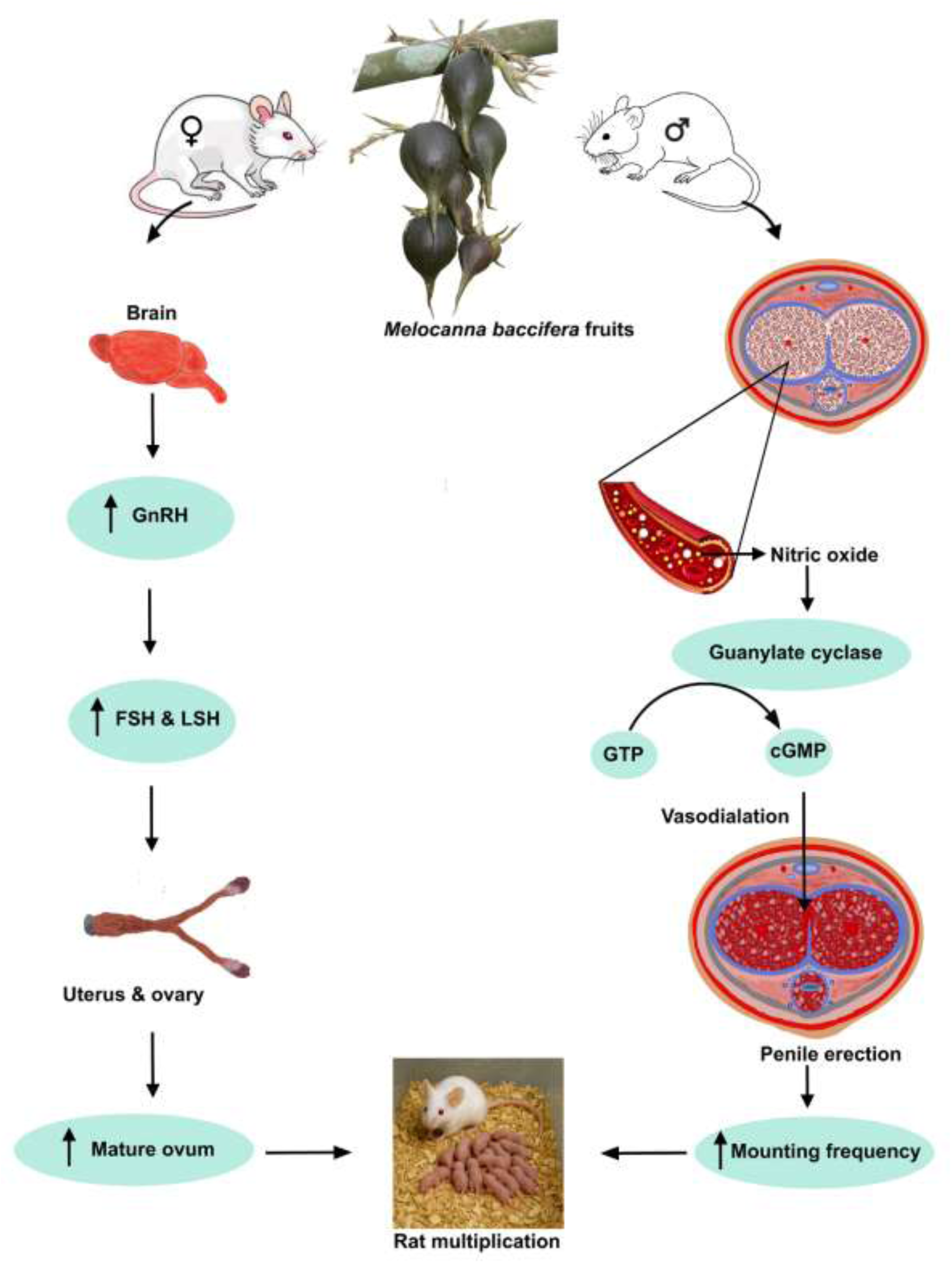
Multiplication mechanism in *M. baccifera* fruit fed rats.

### 4.3. Rat boom in other bamboo species

Rat boom has been reported in a few other bamboo species distributed in India and South America. Mass flowering, production of dry fruits and associated events, known as *Thingtam*, has been reported from *Bambusa tulda* and *Dendrocalamus strictus* from India, but they have not been conclusively associated with large-scale enhancement of rat population (Janzen, 1976; Bhattacharya et al., 2006; Aplin and Lalsiamliana, 2010; Kumawat et al., 2014; Naithani et al., 2025). Similarly, *ratadas* (rodent eruptions or outbreaks) recorded in South America is also not unequivocally linked to bamboo blooms (Jaksic and Lima, 2003). But *Mautam* is a unique ecological phenomenon in which synchronous bamboo flowering (gregarious flowering), mast seeding, and rat boom occur sequentially. Its ecological impact in Mizoram and north east India are well documented over centuries (examples, Ramanayake and Weerawardene, 2003; Shoumatoff, 2008; BAFFACOS, 2009; Aplin and Lalsiamliana, 2010; Wilkins, 2010; Normile, 2010; Koshy et al., 2022). This tri-phasic natural event has significant ecological and public health implications.

Further, the nature and quantity of fruits produced during bamboo flowering play crucial roles in plant-animal (-rat) interactions. For example, the quantitative accounts of fruit production in two species, *Bambusa bambos* (= *B. arundinacea)* and *M. baccifera,* is comparable. In *B. bambos* the fruit is a dry caryopsis which is very small, 0.4-0.8 cm long, ellipsoidal and weighing only 0.012 g. Gadgil and Prasad (1984) reported the seed production by an average *B. bambos* clump as 92 to 103.5 Kg and per hectare production as 10 tonnes. In contrast *M. baccifera* fruits are big, fleshy, baccate, 3.5-11.0 cm long and 2.2-6.0 cm wide with their weight ranging from 47-300 g. During mast seeding a clump of *M. baccifera* produces 37.19 to 456.67 Kg (average 237.18 (± 169.13 Kg) of fruits (Koshy et al., 2022) and 10 to 12 tonnes per hectare (Banik, 2010; Janzen, 1976; Alam, 1995). Due to this surge of fruits in *M. baccifera* the quantity of food available to rats are very high, and being fleshy and sugary they are palatable to them. Here, we infer that the size, abundance, morphology and sugar-amino acid contents of fruits play a significant role in triggering sudden rat flooding (Scheme 1).

Moreover, the quantitative account of rats associated with bamboo fruiting is scarce. The only available data is that of Bamboo Flowering and Famine Combat Scheme implemented in Mizoram where a collection of 1.51 million rats (rat tails) in one year (2006-07) was recorded (BAFFACOS, 2009; Aplin and Lalsiamliana, 2010). In this study, through rat multiplication trials, we demonstrated that just the increase in pups per rat on *M. baccifera* fruit feeding could potentially give rise to c. 81,662 offsprings in three years. Such exponential rise of rats is worrisome as they may cause transmission of many bacterial and viral zoonotic diseases, some of which are detrimental to humans (Fox, 2015). Incidentally, the earliest recorded observation of rat epizootics in India dates back to the 1840s, when British colonial officers reported that Garhwali villagers (in present-day Uttarakhand, northern India) fled to the Himalayan foothills in fear of the ’*Mahamari*’ (epidemic) (Francis and Pearson, 1854; Lynteris, 2022). However, detailed investigations into the rat’s capacity to harbor and transmit diseases began only after the onset of the third plague pandemic in Hong Kong in 1894 (Benedict, 1996; Echenberg, 2007). Furthermore, as demonstrated by Gunn and co-workers (2023) in the case of *Rattus* (black rats), these rodents have been shown to disrupt nutrient pathways, subsequently altering the territorial behaviour of other organisms. In a similar manner, *Mautam*-associated rat outbreaks disrupt nutrient cycles by destroying grain crops, thereby contributing to food shortages and famine conditions (Shoumatoff, 2008; Normile, 2010; Aplin and Lalsiamliana, 2010; Wilkins, 2010). Rat eradication is also considered as a conservation strategy to restore species interactions which can influence populations and communities at higher ecological levels (Gunn et al., 2023). Globally, eradication or control of rats is done by rat-proofing buildings and infrastructures, rat destruction using predators, poisons, viruses, and chemical gases, and hiring skilled rat-catchers (Engelmann, 2021). Such measures involve considerable human resource and massive funding (Aplin and Lalsiamliana, 2010; Fox, 2015, Engelmann, 2021). Our findings may aid reproductive biologists working on targeted interventions in reproductive processes aimed at controlling rat populations.

## 5. CONCLUSIONS

In this study, the enhanced reproductive parameters *viz.*, mounting frequency, serum NO, LH and FSH on feeding *M. baccifera* fruits or the AA mix (Ser-Thr-Glu, three major AAs in *Muli* fruits) are clear indicators towards the high reproductive rates in rats. AA- NO-cGMP/GnRH-FSH-LH mechanism explains these high reproductive rates in *Muli* feeding rats. The sugar mix in *Muli* fruits, primarily sucrose with glucose and fructose, serves as an energy source for reproduction and related biological processes, ultimately triggering *Mautam* approx. twice each century. This is a first study providing an explanation to the rat multiplication mechanism associated with this botanical enigma associated with bamboo flowering. Further, our leads could pave the way for the potential use of amino acids present in *M. baccifera* fruits in the treatment of infertility.

## ACKNOWLEDGEMENTS

We acknowledge Science and Engineering Research Board (SERB), Department of Science and Technology, Government of India for providing financial support (Grant number SR/S0/PS/0015/2011) to this study. This work was also supported by the Plan Funded Project (KSCSTE/JNTBGRI/P-07A) of the Government of Kerala, India.

## AUTHOR CONTRIBUTIONS

S.B., K.C.K. and A.N.S. conceived the idea; K.C.K. and B. Gopakumar provided *M. baccifera* fruits; A.N.S., A.J.J., S.B. and B. Govindan designed and carried out the biological assays; A.N.S., K.C.K. and S.B. wrote the manuscript with inputs from other authors.

## DATA AVAILABILITY STATEMENT

There are no additional data for this paper.

## REFERENCES

1. Alam, M. K. (1995). *Melocanna baccifera* (Roxb.) Kurz., in: Dransfield, S., Widjaja, E. A. (Eds.), Plant Resources of South-East Asia. No. 7: Bamboos. Backhuys, Leiden, pp. 126–129.

2. Aplin, K., Lalsiamliana, J. (2010). Chronicle and impacts of the 2005-09 *mautam* in Mizoram, in: Singleton, G.R, Belmain, S.R., Brown, P.R., Hardy, B. (Eds.), Rodent Outbreaks: Ecology and Impacts. International Rice Research Institute, Manila:, pp. 13–47.

3. Bamboo Flowering and Famine Combat Scheme (BAFFACOS) (2009). Chapter I Performance Review Planning and Programme Implementation Department. Comptroller and Auditor General of India, Government of India. https://cag.gov.in/uploads/download_audit_report/2009/Mizoram_Civil_2009_chap1.p df (accessed 29 May 2025).

4. Banik, R. L. (2010). Biology and Silviculture of Muli Bamboo *Melocanna baccifera* (Roxb.) Kurz. National Mission on Bamboo Applications, New Delhi.

5. Banik, R. L. (2016). Melocanna Trin., in: Silviculture of South Asian Priority Bamboos. Springer Science+Business Media, Singapore, pp. 211–233.

6. Belmain, S. R., Chakma, N., Sarker, N. J., Sarker, S. U., Sarker, S. K., & Kamal, N. Q. (2010). The Chittagong story: studies on ecology of rat floods and bamboo masting, in: Singleton, G. R., Belmain, S. R., Brown, P. R., Hardy, B. (Eds.), Rodent Outbreaks: Ecology and Impacts. International Rice Research Institute, Manila, pp. 49–64.

7. Benedict, C. A. (1996). Bubonic Plague in Nineteenth-Century China. Stanford University Press, California.

8. Bhattacharya, S., Das, M., Bar, R., & Pal, A. (2006). Morphological and molecular characterization of *Bambusa tulda* with a note on flowering. Annals of Botany 98, 529–535. 10.1093/aob/mcl143

9. Bran, D. W. (1995). Glutamate: A major excitatory transmitter in neuroendocrine regulation. Neuroendocrinology, 61, 213–225. 10.1159/000126843

10. Burnett, A. L., Lowenstein, C. J., Bredt, D. S., Chang, T. S., & Snyder, S.H. (1992). Nitric oxide: a physiologic mediator of penile erection. Science, 257, 401–403. doi: 10.1126/science.1378650

11. Burnett, A. L. (1995). Role of nitric oxide in the physiology of erection. Biology of Reproduction, 52, 485–489. 10.1095/biolreprod52.3.485

12. Burnett, A. L. (1997). Nitric oxide in the penis: physiology and pathology. Journal of Urology, 157, 320–324. 10.1016/S0022-5347(01)65369-2

13. Burnett, A. L. (2004). Novel nitric oxide signaling mechanisms regulate the erectile response. International Journal of Impotence Research, 16, S15–S19. 10.1038/sj.ijir.3901209

14. Burnett, A. L. (2006). The role of nitric oxide in erectile dysfunction: implications for medical therapy. The Journal of Clinical Hypertension, 8, 53–62. 10.1111/j.1524-6175.2006.06026.x

15. Burnett, A. L. (2024). Nitric oxide in the penis: still the key erection player? The Journal of Sexual Medicine, 21, 587–588. 10.1093/jsxmed/qdae056

16. Clarkson, J., & Herbison, A. E. (2006). Development of GABA and glutamate signaling at the GnRH neuron in relation to puberty. Molecular and Cellular Endocrinology, *254- 255*, 32-38. 10.1016/j.mce.2006.04.036

17. Diebel, N. D., & Bogdanove, E. M. (1978). Analysis of luteinizing hormone and follicle- stimulating hormone release kinetics during a dynamic secretory event, the postpartum preovulatory surge in the rat, based on quantitative changes in stored and circulating luteinizing hormone and follicle-stimulating hormone and metabolic clearance data for these hormones. Endocrinology, 103, 665–673. 10.1210/endo-103-3-665

18. Dhandapani, K. M., & Brann, D. W. (2000). The role of glutamate and nitric oxide in the reproductive neuroendocrine system. Biochemistry and Cell Biology, 78, 165–179. 10.1139/o00-015

19. Echenberg, M. J. (2007). Plague Ports: The Global Urban Impact of Bubonic Plague, 1894-1901. New York University Press, New York.

20. Engelmann, L. (2021). An epidemic for sale: Observation, modification, and commercial circulation of the Danysz Virus, 1890-1910. Isis, 112, 439–460. doi: 10.1086/715642

21. Farkas, I., Bálint, F., Farkas, E., Vastagh, C., Fekete, C., & Liposits, Z. (2018). Estradiol increases glutamate and GABA neurotransmission into GnRH neurons *via* retrograde no-signaling in proestrous mice during the positive estradiol feedback period. eNeuro, 5(*4*), ENEURO.0057-18.2018. 10.1523/ENEURO.0057-18.2018

22. Fox, J. G. (2015). Diseases transmitted by man’s worst friend: the rat. Microbiology Spectrum, 3(6). 10.1128/microbiolspec.iol5-0015-2015

23. Francis, C. R., & Pearson, F., 1854. “Mahamurree, or Indian plague”. Indian Annals of Medical Science, 2, 609–645.

24. Gadgil, M., & Prasad. S. N. (1984). Ecological determinants of life history evolution of two Indian Bamboo species. Biotropica, 16, 161–172. 10.2307/2388050

25. Gamble, J. S. (1896). Bambuseae of British India. Annals of the Royal Botanic Garden (Calcutta), 7, 133.

26. Govindan, B., Johnson, A., Nair, S. N. A., Gopakumar, B., Mallampalli, K. S. L., Venkataraman, R., Koshy, K. C., & Baby, S. (2016). Nutritional properties of the largest bamboo fruit *Melocanna baccifera* and its ecological significance. Scientific Reports, 6, 26135. 10.1038/srep26135

27. Govindan, B., Johnson, A. J., Gayathri, V., Ramaswamy, V., Koshy, K. C., & Baby, S. (2019). Secondary metabolites from the unique bamboo, *Melocanna baccifera*. Natural Product Research, 33, 122–125. 10.1080/14786419.2018.1434647

28. Gunn, R. L., Benkwitt, C. E., Graham, N. A. J., Hartley, I. R., Algar, A. C., & Keith, S. A. (2023). Terrestrial invasive species alter marine vertebrate behaviour. Nature Ecology & Evolution, 7, 82–91. 10.1038/s41559-022-01931-8

29. Htwe, N. M., Singleton, G. R., Thwe, A. M., & Lwin, Y. Y. (2010). Rodent population outbreaks associated with bamboo flowering in Chin State, Myanmar, in: Singleton, G. R., Belmain, S. R., Brown, P. R., Hardy, B. (Eds.), Rodent Outbreaks: Ecology and Impacts. International Rice Research Institute, Manila, pp. 79-98.

30. Hull, E. M., Lumley, L. A., Matuszewich, L., Dominguez, J., Moses, J., & Lorrain, D. S. 1994. The roles of nitric oxide in sexual function of male rats. Neuropharmacology, 33,1499–1504. 10.1016/0028-3908(94)90054-X

31. Iremonger, K. J., Constantin, S., Liu, X., & Herbison, A. E. (2010). Glutamate regulation of GnRH neuron excitability. Brain Research, 1364, 35–43. 10.1016/j.brainres.2010.08.071

32. Jaksic, F. M., & Lima, M. (2003). Myths and facts on ratadas: Bamboo blooms, rainfall peaks and rodent outbreaks in South America. Austral Ecology, 28, 237–251. 10.1046/j.1442-9993.2003.01271.x

33. Janzen, D. H. (1976). Why bamboos wait so long to flower. *Annual Review of Ecology*, Evolution and Systematics, 7, 347–391. 10.1146/annurev.es.07.110176.002023

34. Jeeva, S., Kiruba, S., Lalhruaitluanga, H., Prasad, M. N. V., & Rao, R. R. (2009). Flowering of *Melocanna baccifera* (Bambusaceae) in northeastern India. Current Science, 96, 1165–1166.

35. John, C. K., & Nadgauda, R. S. (2002). Bamboo flowering and famine. Current Science, 82, 261–262.

36. Koshy, K. C. (2017). Bambusetum at Jawaharlal Nehru Tropical Botanic Garden and Research Institute: State of the art, in: Singh, P., Dash, S. (Eds.), Indian Botanic Gardens: Role in Conservation. Botanical Survey of India, Kolkatta, pp. 361-378.

37. Koshy, K. C. (2023). Germplasm resources of bamboos, in: Das, M., Ma, L., Pal, A., Kole, C. (Eds.), Genetics, Genomics and Breeding of Bamboos. CRC Press, London, pp. 26–115.

38. Koshy, K.C., & Baby, S. (2025). Nutritional and medicinal properties of Melocanna baccifera, in: Pant, M., Husen, A. (Eds.), Exploring Medicinal Bamboos. CRC Press, (in press).

39. Koshy, K. C. (2010). Bamboos at TBGRI. Tropical Botanic Garden and Research Institute, Thiruvananthapuram.

40. Koshy, K. C., Dintu, K. P., & Gopakumar, B. (2010). The enigma of leaf size and plant size in bamboos. Current Science, 99, 1025–1027.

41. Koshy, K. C., Gopakumar, B., Sebastian, A., Nair, A. S., Johnson, A. J., Govindan, B., & Baby, S. (2022). Flower fruit dynamics, visitor-predator patterns and chemical preferences in the tropical bamboo, *Melocanna baccifera*. PLoS One, 17, e0277341. 10.1371/journal.pone.0277341

42. Kumawata, M. M., Singh, K. M., Tripathi, R. S., Riba, T., Singh, S., & Sen, D. (2014). Rodent outbreak in relation to bamboo flowering in north-eastern region of India. Biological Agriculture &. Horticulture, 30, 243–252. 10.1080/01448765.2014.925828

43. Liu, W., Wang, T., Zhao, K., Hanigan, M. D., Lin, X., Hu, Z., Hou, Q., Wang, Y., & Wang, Z. (2024). Effects of individual essential amino acids on growth rates of young rats fed a low-protein diet. Animals, 14, 959. 10.3390/ani14060959

44. Lynteris, C. (2022). The global war against the rat, in: Visual plague: the emergence of epidemic photography. MIT Press, Cambridge. 10.7551/mitpress/14413.001.0001.

45. McClure, F. A. (1966). The Bamboos: A Fresh Perspective. Harward University Press, Cambridge, p. 347.

46. Naithani, H. B., Kant, R., Meena, R. K., Bhandari M. S., & Garg, R. (2025). Gregarious flowering of *Dendrocalamus strictus* in Uttar Pradesh and Maharashtra, India: Compared with historical records. Advances in Bamboo Science, 11, 100167. 10.1016/j.bamboo.2025.100167

47. Normile, D. (2010). Holding back a torrent of rats. Science, 327, 806–807. doi: 10.1126/science.327.5967.806

48. Ramanayake, S. M. S. D., & Weerawardene, T. E. (2003). Flowering in a bamboo, *Melocanna baccifera* (Bambusoideae: Poaceae). Botanical Journal of the Linnean Society, 143, 287–291. 10.1046/j.1095-8339.2003.00216.x

49. Ruiz-Sanchez, E., & Sosa, V. (2015). Origin and evolution of fleshy fruit in woody bamboos. Molecular Phylogenetics and Evolution, 91, 123–134. 10.1016/j.ympev.2015.05.020

50. Sastry, K. V. H., Moudgal, P., Mohan, J., Tyagi, J. S., & Rao, G. S. (2002). Spectrophotometric determination of serum nitrite and nitrate by copper-cadmium alloy. Analytical Biochemistry, 306, 79–82. 10.1006/abio.2002.5676

51. Shoumatoff, A. (2008). Waiting for the plague. http://www.vanityfair.com/news/2007/12/famine200712/ (accessed 06 Apr 2025).

52. Subramoniam, A., Madhavachandran, V., Rajasekharan, S., & Pushpangadan, P. (1997). Aphrodisiac property of *Trichopus zeylanicus* extract in male mice. Journal of Ethnopharmacology, 57, 21–27. 10.1016/S0378-8741(97)00040-8

53. Vorontsova, M. S., Clark, L. G., Dransfield, J., Govaerts, R., & Baker, W. J. (2016). World Checklist of Bamboos and Rattans, INBAR-International Network for Bamboo and Rattan, Beijing & The Board of Trustees of the Royal Botanic Gardens, Kew. https://kew.iro.bl.uk/concern/books/5bee1ffc-a641-4739-9d38-590d7ebb71af.

54. Wang, C., Oh, D.Y., Maiti, K., Kwon, H. B., Cheon, J., Hwang, J.-I., & Seong, J. Y. (2008). Involvement of amino acids flanking Glu7.32 of the gonadotropin-releasing hormone receptor in the selectivity of antagonists. Molecules and Cells, 25, 91–98. 10.1016/S1016-8478(23)17555-8

55. Wilkins, A. (2010). Massive plagues of rats swarm across India every fifty years. https://gizmodo.com/massive-plagues-of-rats-swarm-across-india-every-fifty-5694107/ (accessed 06 Apr 2025).

